# D4: Deep Drug-drug interaction Discovery and Demystification

**DOI:** 10.1101/2020.04.08.032011

**Authors:** Adeeb Noor, Wang Liu-Wei, Ahmed Barnawi, Redhwan Nour, Abdullah A Assiri, Syed Ahmad Chan Bukhari, Robert Hoehndorf

## Abstract

**Motivation:** Drug-drug interactions (DDIs) are complex processes which may depend on many clinical and non-clinical factors. Identifying and distinguishing ways in which drugs interact remains a challenge. To minimize DDIs and to personalize treatment based on accurate stratification of patients, it is crucial that mechanisms of interaction can be identified. Most DDIs are a consequence of metabolic mechanisms of interaction, but DDIs with different mechanisms occur less frequently and are therefore more difficult to identify.

**Results:** We developed a method (D4) for computationally identifying potential DDIs and determining whether they interact based on one of eleven mechanisms of interaction. D4 predicts DDIs and their mechanisms through features that are generated through a deep learning approach from phenotypic and functional knowledge about drugs, their side-effects and targets. Our findings indicate that our method is able to identify known DDIs with high accuracy and that D4 can determine mechanisms of interaction. We also identify numerous novel and potential DDIs for each mechanism of interaction and evaluate our predictions using DDIs from adverse event reporting systems.

**Availability:** https://github.com/bio-ontology-research-group/D4

## 1 Introduction

Concern about drug-drug interactions (DDI) in patients receiving multi-drug therapy has risen in recent years (Qato *et al*., 2008) because such patients may be at high risk. DDIs were found in the U.S to be linked with 0.054% of visits to the ER, 0.57% of admissions to hospital and 0.12% of re-hospitalizations (Becker *et al*., 2007). It is expected that the impact of DDIs on patients’ health will significantly increase as a result of a rise in the number of drugs being prescribed to each individual patient (Percha and Altman, 2013). Early identification of DDIs is challenging due to a number of factors including the lack of information on DDIs prior to a new drug being released to the market (Reis and Cassiani, 2010; LePendu *et al*., 2013). The minimal information gained from clinical trials conducted before a new drug is approved is generally not sufficient to thoroughly test interactions with other medications, in particular where these are not administered to patients participating in the study. Additionally, a narrow focus on single mechanisms of interaction (Huitema *et al*., 2001) constitute another factor that increases the challenge to identify and prevent DDIs.

There are now a number of computational methods that are able to identify different mechanisms of interaction in DDIs (Berger and Iyengar, 2011), focusing mainly on the pharmacodynamic interactions (drug– target, therapeutic and adverse drug effect) or pharmacokinetic interactions (drug–protein). For those interactions, measures of similarity among drugs and mechanistic information (Ferdousi *et al*., 2017), Semantic Web Technologies and Linked Data in the life sciences (Kamdar and Musen, 2017), web data (Fokoue *et al*., 2016), textual (Abdelaziz *et al*., 2017; Tari *et al*., 2010), and interaction networks (Bai and Abernethy, 2013; Park *et al*., 2015) have been developed and successfully applied to predict DDIs and ways in which drugs interact. Moreover, features that have been used to identify and better understand DDIs and mechanisms of interaction include molecular structure (Vilar *et al*., 2012), interaction profile fingerprints (Vilar *et al*., 2013), phenotypic, therapeutic, chemical, and genomic properties (Cheng and Zhao, 2014; Ferdousi *et al*., 2017), model organism phenotypes (Hoehndorf *et al*., 2013), multiple interaction mechanisms (Noor *et al*., 2016), and drug and protein properties (Kamdar and Musen, 2017).

These methods can predict and identify different mechanisms of interaction in DDIs. The majority of computational methods focuses on single mechanisms of interaction (mainly interactions through drug metabolic pathways) (Preissner *et al*., 2009; Tari *et al*., 2010; Leone *et al*., 2010; Fournier *et al*., 2014). However, several mechanisms for DDIs exist and some drugs interact through multiple mechanisms, such as the interactions between statins and cyclosporine occurring through the metabolism (CYP3A4) and transport (P-glycoprotein) pathways (Holtzman *et al*., 2006). One challenge in predicting or inferring DDIs across multiple mechanisms of interaction are the variety of features that are required for prioritizing the different mechanisms. Another limitation can be the use of a single training or evaluation dataset which may be biased towards particular types of interactions. For example, methods that rely on DDIs in the DrugBank database (Wishart *et al*., 2017) for training or testing may perform well on the types of interactions in that database but may not generalize to other datasets that include different types of interactions (Peters *et al*., 2015; Grizzle *et al*., 2019).

A large number of information that may be used to predict DDIs across multiple mechanisms is available in biological knowledge bases and enriched with background knowledge through biomedical ontologies (Hoehndorf *et al*., 2015). Recent deep learning methods have been used successfully in multiple domains (LeCun *et al*., 2015), including prediction of DDIs (Ryu *et al*., 2018) and on structured knowledge bases and ontologies (Alshahrani and Hoehndorf, 2018; Smaili *et al*., 2018). A key advantage of relying on structured knowledge is the ability to qualitatively identify and interpret some prediction results.

We have developed a method for predicting DDIs together with their mechanisms of interaction. Our method relies on a large set of different DDI resources for training and evaluation, and a knowledge-based algorithm that determines the likely mechanism through which a known DDI arises. We generate features for drugs based on structured knowledge and ontology-based annotations of drugs representing their side effects and the functions of their targets, and we use these features as input to a neural network classifier. We demonstrate that our method accurately identifies known DDIs together with the mechanism of interaction, and we show how our method can be used to discover new potential DDIs.

## 2 Results

### 2.1 Representation learning and feature generation

We use information about drug side effects, functions of drug protein targets, and the mechanisms of interaction associated with known DDIs in order to predict novel DDIs and their mechanisms. In addition to this information, we also use background information about the relationship between phenotypes, biological functions and processes, and pharmacological data in the form of logical axioms and natural language definitions represented in biomedical ontologies.

We represent drugs with their associated side effects (phenotypes) from the SIDER database (Kuhn *et al*., 2015) and with the functions and phenotypes associated with their protein targets using the set of targets in DrugBank (Wishart *et al*., 2017). We use the phenotype annotations of side effects from the Human Phenotype Ontology (HPO) database (Köhler *et al*., 2018) and function annotations from the Gene Ontology (GO) database (Ashburner *et al*., 2000; Consortium, 2016). We associate a total of 848 drugs with side effects and 827 drugs with the GO functions of their targets.

Each of these features is based on biomedical ontologies, in particular the GO and HPO. Therefore, we also use the axioms in these ontologies as background knowledge during feature generation. We use the OPA2Vec tool (Smaili *et al*., 2018) to encode drugs, their ontology-based associations, and the ontology axioms as *n*-dimensional feature vectors. OPA2Vec treats the axioms and meta-data in ontologies, and the ontology-based annotations of the drugs, as a corpus and uses Word2Vec to encode the drug identifiers, their annotations and the ontologies as features vectors (Smaili *et al*., 2018). To determine the effect of the different features (annotations and ontologies) on the performance of our DDI prediction models, we first generate features separately for each type of association. Due to the different coverage of drugs with side effects and functions of their targets, we also considered the intersections between phenotype and functional annotations as well as the union. We use these feature vectors as input to train a neural network to prediction DDIs after using a rule-based approach to distinguish between their mechanisms of interaction. Figure 1 shows the workflow of our prediction method.

**Fig. 1:**
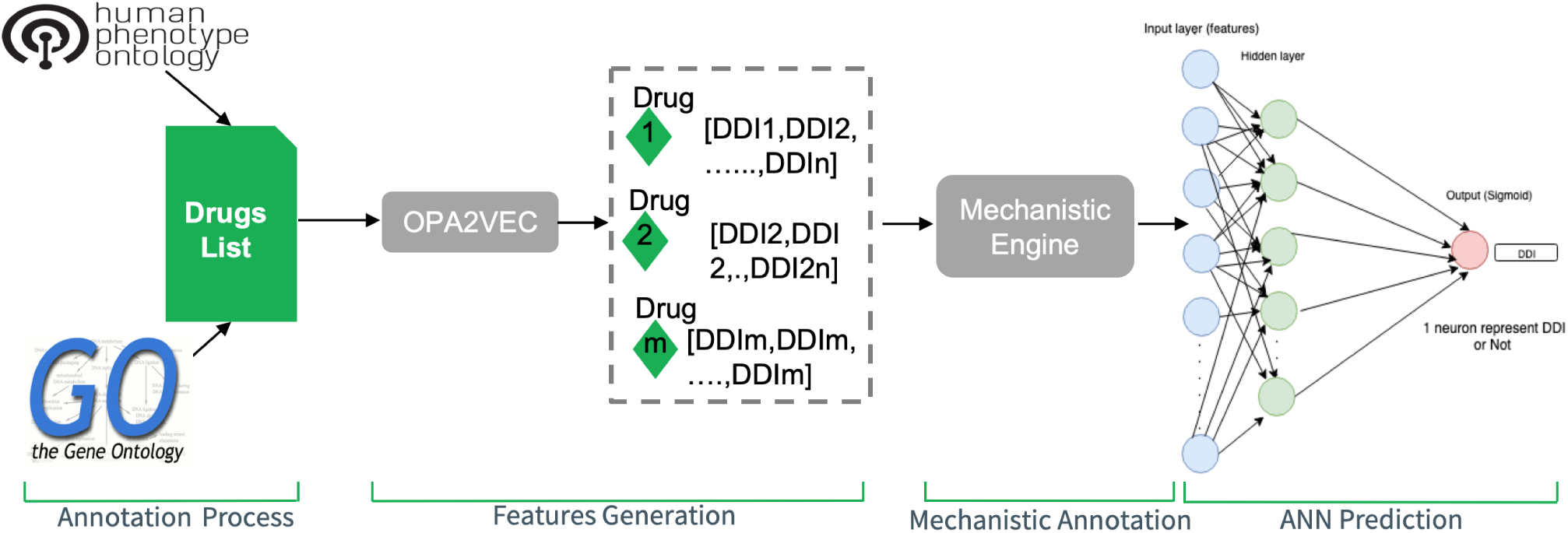
D4 workflow. Starting with drug annotations from HPO and GO, vectors were generated with OPA2Vec followed by annotation of DDIs with their associated mechanism of action. The prediction process done by training ANN in order to which mechanism of action each DDI belong to.

### 2.2 Annotation with DDI mechanisms

We generate a model based on supervised training data using only DDIs that have been observed in a clinical context (in contrast to predicted DDIs). We obtain DDIs from the Potential Drug-Drug Interaction (PDDI) dataset (Ayvaz *et al*., 2015), an aggregation of 17 datasets of which 9 have clinical evidences, with a resulting 39,815 pairs of DDIs. As we intend to distinguish between different mechanisms of interaction, we first annotate these DDIs with their interaction mechanisms.

We broadly distinguish between mechanisms of DDIs due to pharmacokinetic, pharmacodynamic, multiple-pathway, and pharmacogenetic interactions. Specifically, we utilize a rule-based inference engine for drug-drug interaction discovery and demystification (D3) (Noor *et al*., 2016) to annotate known DDIs with their mechanisms of interaction. D3 applies rules on a knowledge graph to distinguish between five pharmacokinetic mechanisms of interaction: protein binding, metabolic induction, metabolic inhibition, transporter induction, and transporter inhibition. D3 obtains information about these mechanisms from DrugBank (Wishart *et al*., 2017) which characterizes DDIs on the protein level. Pharmacodynamic mechanisms of interaction include both competitive interactions (drugs sharing the same biological targets) and additive interactions (drugs sharing the same side effects, same mechanisms of interaction, or same indications). The information about pharmacodynamic interactions in D3 is obtained from DrugBank (Wishart *et al*., 2017), from SIDER for side effects (Kuhn *et al*., 2015), and the National Drug File Reference Terminology (NDF-RT) (Brown *et al*., 2004) for competitive action and indication. In addition, D3 obtains information about multi-pathway interactions (drugs sharing at least one metabolism and transport) from DrugBank, while the pharmacogenetic information is from PharmGKB (Klein *et al*., 2001). We also add a “generic” mechanism of interaction, i.e., a DDI with no specific mechanism of interaction. The results are 11 mechanisms of interaction and one class for “generic” DDIs.

### 2.3 Performance in predicting interactions using phenotypic and functional knowledge

We then use pairs of feature vectors as input for training an Artificial Neural Network (ANN) to predict DDI interactions. We applied leave-one-drug-out cross-validation in which one drug is held out for validation while all the other drugs and their interactions were used as training data. This method is different from splitting pairs of drugs and intended to avoid training a model that primarily predicts interactions based on other interactions seen during training. We evaluate the resulting model by ranking, for each drug, all other drugs based on the output of the ANN sigmoid classification score. We train separate models for each mechanism of interaction, and Figure 2 shows the receiver operating characteristic (ROC) curves for pharmacokinetic mechanism, pharmacodynamic mechanism, multi-pathway mechanism, and pharmacogenetic mechanism (complete results for all interaction mechanisms are provided as Supplementary Figure 1. Table 1 provides a summary of the performance for the prediction of each mechanism.

**Table 1.** Mechanisms of interaction used for phenotypic and functional predictions along with the pharmacology levels and the ROCAUC for all features. The features are Human Phenotype Ontology (HP) associations and Gene Ontology (GO) associations, their intersection and their union.

| Mechanism of Interaction | Pharmacology Level | Features AUC score |  |  |  |
| --- | --- | --- | --- | --- | --- |
|  |  | HPO | GO | HPO∩GO | HPO∪GO |
| Protein-binding | Pharmacokinetic | 0.698 | 0.730 | 0.702 | 0.781 |
| Metabolic-induction | Pharmacokinetic | 0.762 | 0.816 | 0.792 | 0.856 |
| Metabolic-inhibition | Pharmacokinetic | 0.727 | 0.785 | 0.740 | 0.820 |
| Transporter-induction | Pharmacokinetic | 0.824 | 0.829 | 0.842 | 0.894 |
| Transporter-inhibition | Pharmacokinetic | 0.788 | 0.803 | 0.780 | 0.843 |
| Biological-targets | Pharmacodynamic | 0.890 | 0.930 | 0.904 | 0.947 |
| Side-effects | Pharmacodynamic | 0.668 | 0.724 | 0.677 | 0.747 |
| Mechanism-of-action | Pharmacodynamic | 0.755 | 0.851 | 0.787 | 0.855 |
| Indications | Pharmacodynamic | 0.887 | 0.929 | 0.905 | 0.947 |
| Metabolism-Transporter | Multi-pathway | 0.817 | 0.833 | 0.811 | 0.873 |
| SNPs | Pharmacogenetic | 0.833 | 0.846 | 0.829 | 0.878 |

**Fig. 2:**
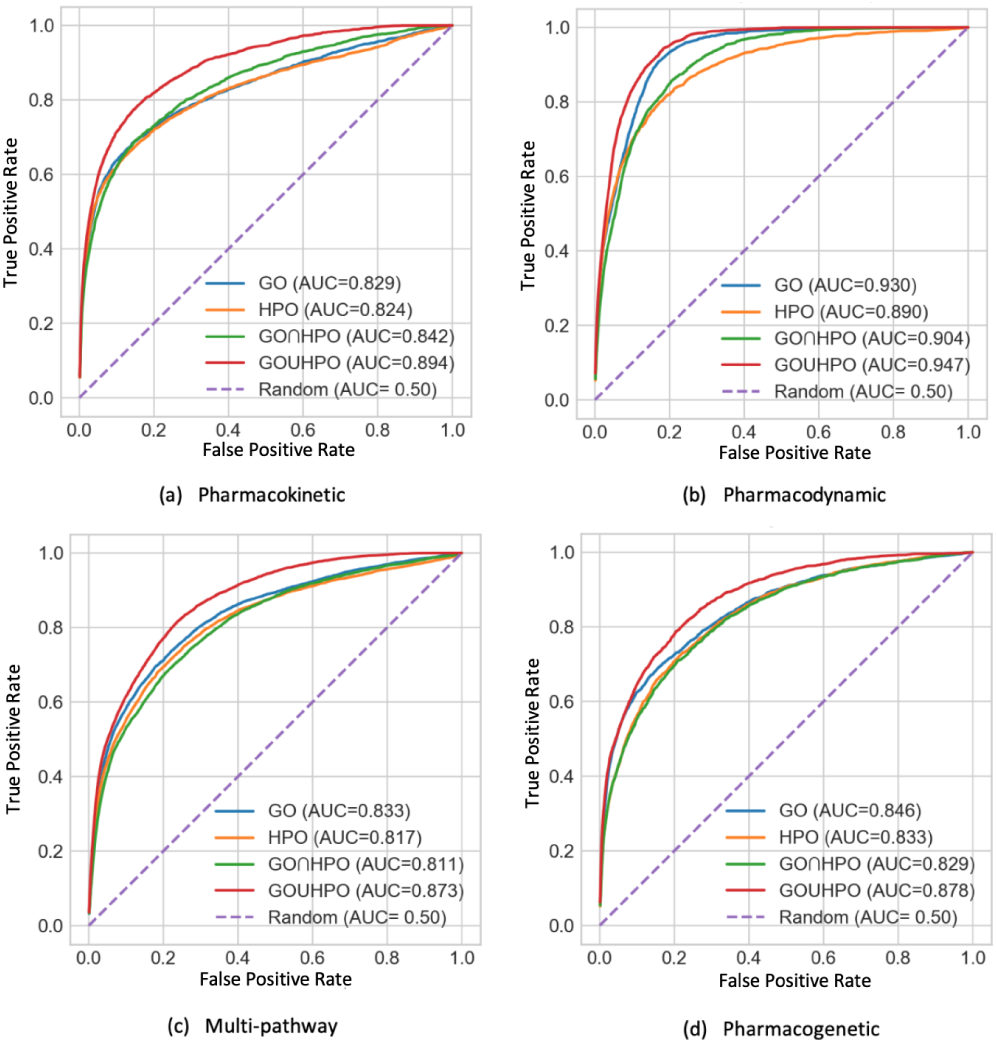
The ROC curves (micro-average over all drugs) for DDI prediction for four different mechanisms: pharmacokinetic, pharmacodynamic, multi-pathway, and pharmacogenetic interactions.

Our evaluation procedure is internal and may not translate to other sets of drugs or DDIs. To provide an external evaluation of our method, we use the TWOSIDES dataset, a dataset of potential DDIs that have been derived statistically from adverse event reporting systems and electronic health records (Tatonetti *et al*., 2012). TWOSIDES provides statistical evidence for potential DDIs and associates each candidate DDI with a *p*-value. If our model generalizes to datasets such as TWOSIDES, we expect that high-confidence DDIs in TWOSIDES are also predicted as DDIs by our method. For the purpose of our evaluation, we consider any DDI in TWOSIDES associated with a *p*-value less than 0.05 as a positive and compute the overall recall and precision using these positives obtained from TWOSIDES. We also remove duplicates DDI between TWOSIDES and our training before testing.

Table 2 shows the results of our comparison. The highest recall across all mechanisms of interaction is 0.58 for the metabolic inhibition mechanism while the lowest recall is 0.26 for DDIs sharing the same indication. Our method achieves 0.22 as the highest precision for DDIs sharing the same SNPs and the lowest precision is 0.09 for sharing the same mechanism of action. When comparing against potential DDIs in TWOSIDES, the ROCAUC of our method is 0.99 for DDIs sharing the same mechanism of action, while the lowest ROCAUC (0.854) is for DDIs with the same side effects (see Supplementary Figure 2). All of our predictions together with the predicted mechanism and the associated TWOSIDES confidence can be found at https://bio2vec.cbrc.kaust.edu.sa/data/D4/predictions/.

**Table 2.** Recall and precision for D4 predictions when compared to TWOSIDES associations with a *p*-value less than 0.05.

| Mechanism of Interaction | TWOSIDES |  |
| --- | --- | --- |
|  | Recall | Precision |
| Protein-binding | 0.471 | 0.137 |
| Metabolic-induction | 0.581 | 0.115 |
| Metabolic-inhibition | 0.585 | 0.124 |
| Transporter-induction | 0.434 | 0.203 |
| Transporter-inhibition | 0.509 | 0.154 |
| Biological-targets | 0.268 | 0.181 |
| Side-effects | 0.551 | 0.138 |
| Mechanism-of-action | 0.559 | 0.096 |
| Indications | 0.268 | 0.181 |
| Metabolism-Transporter | 0.444 | 0.177 |
| SNPs | 0.464 | 0.225 |
| DDIs | 0.534 | 0.149 |

## 3 Discussion

Effective early detection of DDIs has been a desirable goal for pharmaceutical companies and clinicians to avoid serious health complications for patients. A variety of studies have been done for discovering DDIs based on clinical and computational approaches, for example through mining DDIs from scientific literature (Tari *et al*., 2010; Percha *et al*., 2012), Adverse Event Reporting Systems (Ibrahim *et al*., 2016), and the Electronic Health Record (Pathak *et al*., 2013). Clinical studies are conducted to determine suspected interactions (Wienkers and Heath, 2005), yet their investigative processes are slow and often consider only a small numbers of drugs and targets (Bjornsson *et al*., 2003) suspected to result in interactions. Our method can be used to suggest potential interactions to consider in clinical studies so as to improve their efficiency and detect interactions that may not be obvious. Suggesting the mechanisms in addition to the mere presence of an interaction can further be used to design targeted studies.

We developed the D4 method that predicts DDIs and identifies the role of mechanisms of interaction by using the phenotypic, functional, and mechanistic features of drugs. Our method is based on incorporating a large volume of biological background information in the form of ontologies and annotations of drugs. This background knowledge is encoded using a machine learning method that combines ontology axioms with structured annotations of drugs and their targets. While there is a large number of methods to predict DDIs (Ryu *et al*., 2018; Zhou *et al*., 2018; Wang *et al*., 2019; Sun *et al*., 2019), D4 specifically predicts the mechanisms of the interaction, focusing on 11 common mechanisms. We have validated our method both on the training data we used as well as on an external dataset obtained from mining adverse event reporting systems and could demonstrate the our method can accurately predict DDIs.

D4 has some limitations that should be addressed in future work. First, for a new DDI that occurs due to chemical structure or interaction, D4 method will currently not be able to identify any interactions because D4 does not consider the structural properties of the chemical; instead, D4 relies only on publicly available, semantically coded qualitative information about drugs. One possibility to address this issue is combine D4 with DDI prediction methods that utilize structure (Ryu *et al*., 2018). Second, DDIs are complex processes that are commonly the result of multiple factors, and D4 will only predict individual types of interactions and not consider their potential interactions. Incorporating additional interactions, including protein-protein interactions and metabolic interactions may further improve the utility of D4.

## 4 Methods

### 4.1 Data sources

To generate drug–phenotype associations, we use the SIDER database (Kuhn *et al*., 2015) downloaded on November 10, 2018 (http://sideeffects.embl.de/media/download/meddra_all_se.tsv.gz). We map the side-effects, represented by their UMLS identifiers, to their respective HPO terms using a groovy script using owl-api.

As a result we obtain 62,777 non-duplicate drug–HPO associations. To associate drugs with functions from the Gene Ontology, we use the drug-target associations (https://www.drugbank.ca/ releases/5-1-2/downloads/target-approved-polypeptide-sequences) from DrugBank (Wishart *et al*., 2017), downloaded on November 20, 2018. To annotate targets with functional information, we use the GO annotations from the GO database (Ashburner *et al*., 2000), downloaded on November 25, 2018. In total, we obtain 78,137 unique drugs–GO associations.

As resource of DDIs for training, we downloaded the PDDI dataset (Ayvaz *et al*., 2015) (https://github.com/dbmi-pitt/public-PDDI-analysis/tree/master/) on November 10, 2018, and we use a selection of more “conservative” datasets: NDF-RT, Crediblemeds, ONC-HighPriority, ONC-Non-interruptive, OSCAR, HIV, HEP, and FRENSH. We combine and remove overlap between these resources and then map them to STITCH identifiers using their Anatomical Therapeutic Chemical (ATC) identifiers. As a consequence, we obtain 39,815 unique pairs of DDIs.

We represent phenotypes using the cross-species PhenomeNET ontology (Hoehndorf *et al*., 2011), obtained from the AberOWL ontology repository on November 10, 2018, and we represent functions and cellular locations using the Gene Ontology obtained from the Gene Ontology website (http://www.geneontology.org/page/download-ontology) on November 10, 2018.

We use the TWOSIDES database (Tatonetti *et al*., 2012), downloaded on November 20, 2018, for additional validation. TWOSIDES provides a *p*-value for each potential DDI, and we use a threshold of ≤ 0.05 to consider interactions in TWOSIDES as positive.

### 4.2 Feature generation through embeddings

We use the latest version of OPA2Vec (Smaili *et al*., 2018) downloaded from (https://github.com/bio-ontology-research-group/opa2vec) on December 5, 2018 to generate feature vectors from ontologies and their annotations. OPA2Vec generates a corpus characterizing drugs based on the drug entity, its annotations, class axioms, and metadata from the ontologies. To generate drug embeddings based on phenotypes (side effects), drugs are gathered from SIDER database (drugs-side-effects associations); drugs that do not exist in our gold-standard are removed, leading to 849 drugs. OPA2Vec then generates embeddings for the list of drugs (849 entities) using the following inputs: 1) drugs associations with phenotypes (62,777 associations), and the PhenomeNET.owl ontology file, and the drug identifiers representing types of drugs.

For the drugs embeddings based on the GO functions of the drugs’ targets we collected drugs from DrugBank and removed drug identifiers that do not exist in our evaluate set, resulting in 827 drugs. Then these 827 drug representations with 78,137 drug–GO associations, together with the OWL version of GO, were used as inputs for OPA2Vec to generated the drug embeddings. We considered also the union and intersections when generating the embeddings because of the different coverage of drugs with side effects and functions of their targets. The union and interactions between HPO and GO was computed on the list of drugs we conclude and their associations; PhenomeNET.owl was used as ontology as it includes the GO. Therefore, four different embeddings representing drugs were generated.

For all four embeddings, we used the following OPA2Vec parameters: the skipgram model with window size 5 and a minimum count of 2, and an embedding size of 200. All of the embeddings we generated can be found on https://bio2vec.cbrc.kaust.edu.sa/data/D4/embeddings/.

### 4.3 D4 mechanistic inferential engine

For predicting mechanisms of interaction, we used the D3 system (Noor *et al*., 2016) which is able to suggest mechanisms for an observed DDI. To assure the completeness and update of the information, we updated the knowledge graph to the last update from DrugBank (https://www.drugbank.ca/releases/5-1-2/downloads/all-full-database), SIDER (http://sideeffects.embl.de/media/download/meddra_all_se.tsv.gz), NDF-RT from Unified Medical Language System (UMLS) (Bodenreider, 2004), and PharmGKB (Hewett *et al*., 2002) (https://www.pharmgkb.org/downloads) sources.

In addition, we added a new pharmcodynamic inference based on the following rule: if two drugs treat the same disease they may interact. We also separate the additive pharmacodynamic mechanisms used in the previous work on the D3 system into two inferences: 1) same mechanisms of interaction and 2) same side-effects. These modifications ensure that inferred DDI mechanisms are mutually exclusive. Supplementary Table 1 illustrates the condition of each mechanism of interaction along with sources of information.

### 4.4 D4 supervised artificial neural network model

We trained a feed-forward neural network to predict whether two drugs interact based on the features vectors (generated by OPA2Vec) for both drugs. To optimize hyperparameters of the neural network classifier, we optimized hyperparameters using Hyperas (https://github.com/maxpumperla/hyperas). The neural network architecture is provided in Supplementary Table 2.

The input to the neural network classifier is a vector of dimension 400, representing two features vectors generated for two drugs. The output is a sigmoid classifier indicating whether a DDI exists, or what mechanism of interaction underlies a DDI. In our evaluation, we fix one drug and predict potential interactions for all other drugs; we then rank the prediction scores from the sigmoid classifier for all potential interaction partners. We compute the TPR curve by plotting the true positives rate as a function of the false positive rate; statistics of true and false positives are computed per drug and plots show the results based on the micro average (per drug).

