## Supplementary material for "D4: Deep Drug-drug interaction Discovery and Demystification": Table1, Table2, Fig1, Fig2

*Bioinformatics*
doi.10.1093/bioinformatics/xxxxxx
Advance Access Publication Date: Day Month Year
Manuscript Category

OXFORD

Subject Section

### Supplementary materials to: D4: Deep Drug-drug interaction Discovery and Demystification

**Adeeb Noor [1,*], Wang Liu-Wei [2], Ahmed Barnawi [1], Redhwan Nour [3], Abdullah A Assiri [4], Syed Ahmad Chan Bukhari [5] and Robert Hoehndorf [2,*]**

[1]Department of Information Technology, Faculty of Computing and Information Technology, King Abdulaziz University, Jeddah 80221, Saudi Arabia
[2]King Abdullah University of Science and Technology (KAUST), Computational Bioscience Research Center (CBRC), Computer, Electrical & Mathematical Sciences and Engineering (CEMSE) Division, Thuwal 23955, Saudi Arabia
[3]Department of Computer Science, Taibah University, Medina 42353, Saudi Arabia
[4]Department of Clinical Pharmacy, College of Pharmacy, King Khalid University. Abha, 62529, Saudi Arabia.
[5]Division of Computer Science, Mathematics and Science Lesley H. and William L., Collins College of Professional Studies, St. John's University, New York, NY, USA

*To whom correspondence should be addressed.

Associate Editor: XXXXXXX

Received on XXXXX; revised on XXXXX; accepted on XXXXX

**Abstract**

Table 1. **Mechanisms of interaction used by D4.** Explanation for the condition of interaction along with source of information is provided

| Mechanism of Interaction | Pharmacology Level | Conditions of Interaction | Source of Information |
|---|---|---|---|
| Protein-binding | Pharmacokinetic | drugs bind to same protein > 70% | DrugBank |
| Metabolic-induction | Pharmacokinetic | induction of protein | DrugBank |
| Metabolic-inhibition | Pharmacokinetic | inhibition of protein | DrugBank |
| Transporter-induction | Pharmacokinetic | induction of protein | DrugBank |
| Transporter-inhibition | Pharmacokinetic | inhibition of protein | DrugBank |
| Biological-targets | Pharmacodynamic | drugs share at least one biological target | DrugBank |
| Side-effects | Pharmacodynamic | drugs share at least one side-effect | SIDER |
| Mechanism-of-action | Pharmacodynamic | drugs share at least one mechanism of action | NDF-RT |
| Indications | Pharmacodynamic | drugs treat same disease | NDF-RT |
| Metabolism-Transporter | Multi-pathway | drugs share at least one protein at metabolism and transporter | DrugBank |
| SNPs | Pharmacogenetic | drugs share at least one single-nucleotide polymorphism | PharmGKB |

Table 2. **Best performing model chosen hyper-parameters.** We used hyperas for all 11 mechanisms of interaction in addition to the binary DDI.

| Mechanism of Interaction | Number of Layers | Activation Function | Learning Rate | Optimizer |
|---|---|---|---|---|
| Protein-binding | 16 | ReLU | 0.020 | Adam |
| Metabolic-induction | 256 | ReLU | 0.112 | Adam |
| Metabolic-inhibition | 256 | ReLU | 0.112 | Adam |
| Transporter-induction | 256 | ReLU | 0.144 | Adam |
| Transporter-inhibition | 128 | ReLU | 0.089 | Adam |
| Biological-targets | 256 | ReLU | 0.042 | Adam |
| Side-effects | 256 | ReLU | 0.174 | Adam |
| Mechanism-of-action | 256 | ReLU | 0.194 | Adam |
| Indications | 256 | ReLU | 0.020 | Adam |
| Multi-pathway | 256 | ReLU | 0.112 | Adam |
| SNPs | 256 | ReLU | 0.324 | Adam |
| DDI | 256 | ReLU | 0.185 | Adam |

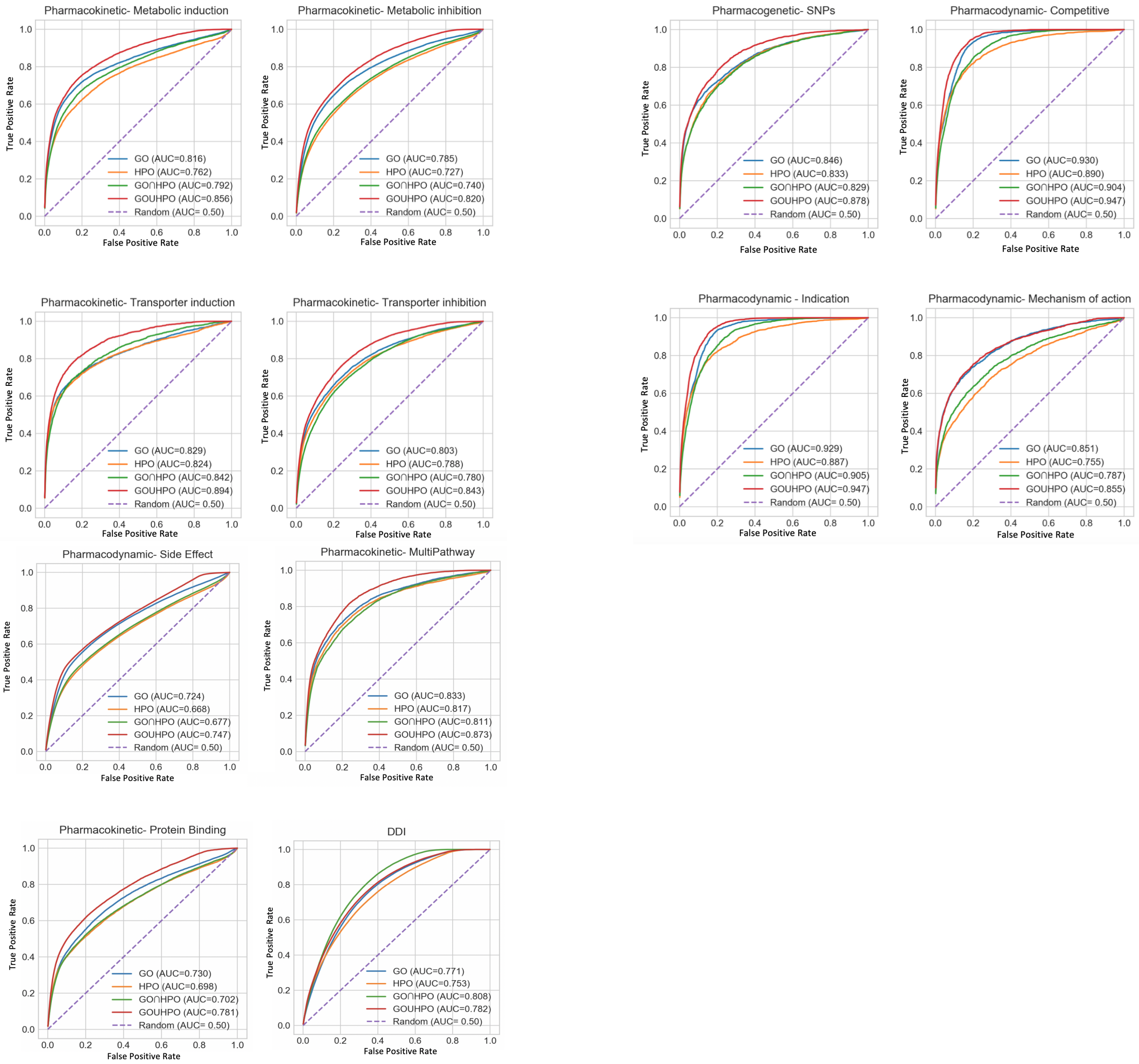


Fig. 1: AUC curves for 12 DDIs model by D4

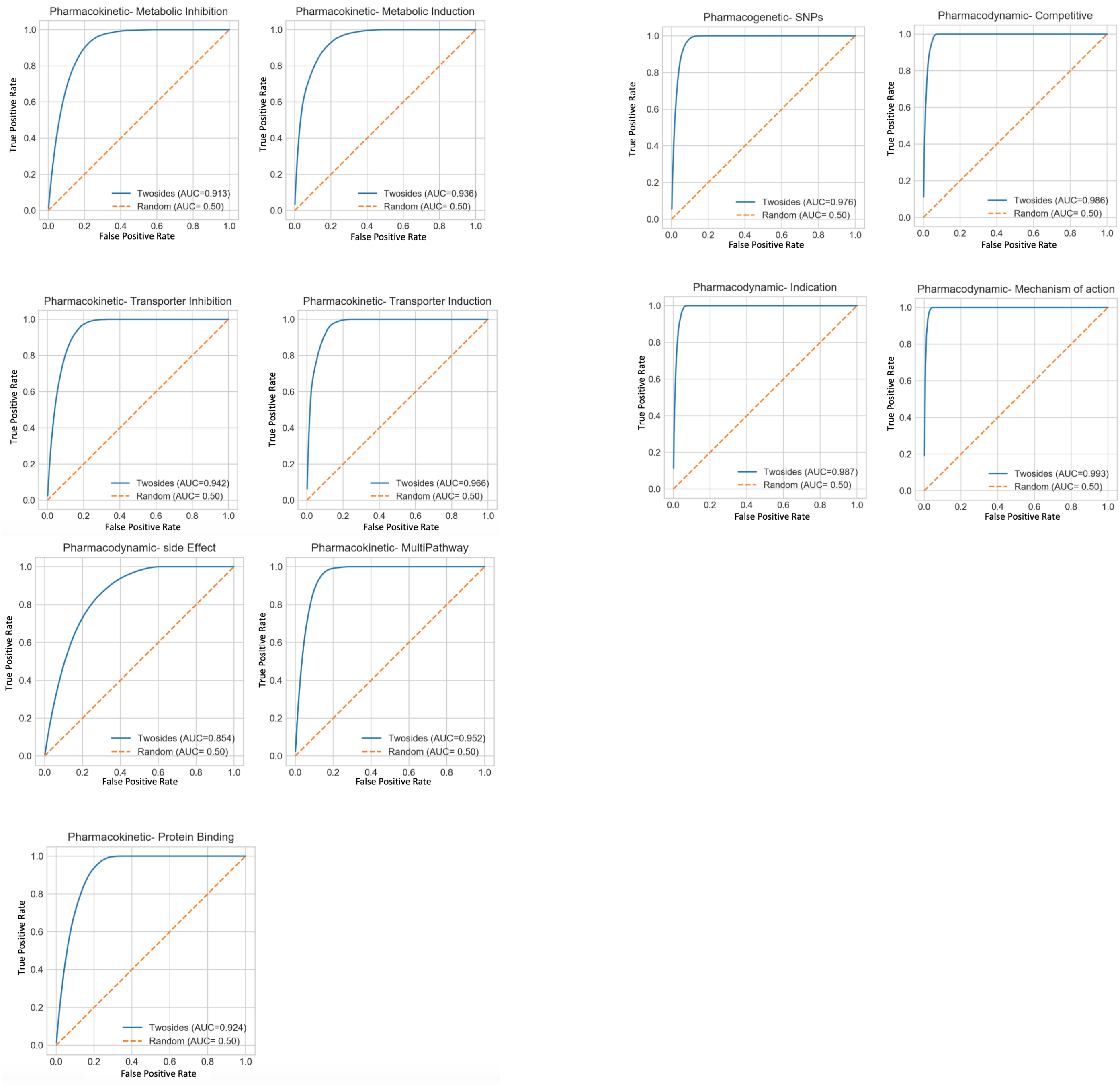


Fig. 2: AUC curves for 11 DDIs model by D4 within TWOSIDES dataset
